# Intertrial Variability in Human Corticospinal Activity during Grasp Force Planning

**DOI:** 10.1101/676833

**Authors:** Nishant Rao, Pranav J. Parikh

**Author notes:** **Corresponding Author:** Pranav J. Parikh, Ph.D., Department of Health and Human Performance, 3875 Holman Street, suite 104R GAR, University of Houston, Houston, TX 77204, USA.

## Abstract

Neuronal firing rate variability during planning has been found to contribute to trial-to-trial variability in primate behavior. However, in humans, whether planning related mechanisms contribute to trial-to-trial behavioral variability remains unknown. We investigated the time-course of trial-to-trial variability in corticospinal excitability (CSE) using transcranial magnetic stimulation (TMS) while subjects planned to perform a self-paced reach-to-grasp task. We hypothesized that CSE variability will be modulated during task planning and that such a modulation would explain trial-to-trial behavioral variability. Able-bodied individuals were visually cued to plan their grip force before exertion of either 30% or 5% of maximum force on an object. TMS was delivered at different time points following a cue that instructed the force level. We first modeled the relation between CSE magnitude and its variability at rest (n=12) to study the component of CSE variability during task planning that was not related to changes in CSE magnitude (n=12). We found an increase in CSE variability during task planning at 30% but not at 5% of force. This effect was temporally dissociated from the decrease in CSE magnitude. Importantly, the increase in CSE variability during planning explained 64% of inter-individual differences in time to peak force rate trial-to-trial variability. These results were found to be repeatable across studies and robust to different analysis methods. Our findings suggest that the planning-related mechanisms underlying modulation in CSE variability and CSE magnitude are distinct. Notably, the extent of modulation in planning-related variability in corticospinal system within individuals may explain their trial-to-trial behavioral variability.

## INTRODUCTION

Trial-to-trial variability is an inherent feature of motor behavior (Stein et al., 2005; Faisal et al., 2008). Intertrial variability in motor output reflects the presence of stochastic noise in the sensorimotor system and may interfere with one’s ability to perform a given movement consistently (Harris and Wolpert, 1998; Slifkin and Newell, 1999; Stein et al., 2005; Faisal et al., 2008). Another perspective suggests that the intertrial motor output variability provides the sensorimotor system an ability to explore the motor workspace for optimizing motor learning (Tumer and Brainard, 2007; Wu et al., 2014).

Several studies have found central correlates of variability in kinematic or kinetic features of motor output (Osborne et al., 2005; Churchland et al., 2006a; Fox et al., 2007; Hohl et al., 2013; Lisberger and Medina, 2015; Haar et al., 2017). In monkeys, variable activity of sensory neuronal populations within extrastriate MT region explained variability in execution of smooth-pursuit eye movement (Hohl et al., 2013). Firing rates of neurons within primate primary motor (M1) and premotor cortices during movement preparation explained intertrial variability in peak reach velocity (Churchland et al., 2006a). In humans, variation in fMRI responses within inferior parietal lobule observed during motor execution has been shown to explain differences in intertrial variability in reach kinematics across individuals (Haar et al., 2017). However, in humans, the role of planning related central mechanisms to variability in motor output remains to be known.

Motor evoked potentials (MEP) elicited non-invasively using transcranial magnetic stimulation (TMS) can provide information regarding the neural mechanisms at cortical, subcortical, and spinal levels, i.e. corticospinal excitability (CSE), during various phases of a task (Bestmann and Krakauer, 2015). The modulation in intertrial variability in CSE assessed during movement preparation has been shown to encode value-based decision-making processes and differentiate fast versus slow reaction time responses (Klein-Flugge et al., 2013). These findings suggest that the temporal unfolding of CSE variability may provide information regarding planning-related neural processes. In the current study, we investigated the time course of CSE variability while able-bodied individuals prepared to perform a self-paced, isometric grip force production task and studied whether the modulation in CSE variability explained differences in trial-to-trial variability in the application of grip force across individuals. Subjects were instructed to first reach for an instrumented object, grasp it, and apply grip force. They were cued to exert either 30% or 5% of the maximal pinch force during the task. We delivered TMS pulses over M1 at different time points during the planning phase of the task to assess the temporal unfolding of CSE variability. Intertrial variability in CSE assessed in this manner may be related to changes in MEP amplitude, a phenomenon that has been studied before (Stein et al., 2005; Darling et al., 2006; Faisal et al., 2008; Bestmann and Krakauer, 2015). Therefore, we modeled a relation between CSE variability and its amplitude in absence of a task during a separate session (Darling et al., 2006; Klein-Flugge et al., 2013). This allowed us to study the component of CSE variability that was beyond the intrinsic changes in CSE magnitude. We hypothesized that the planning-related CSE variability would be modulated while preparing to perform the force production task. Furthermore, we expect that individuals with greater modulation in CSE variability would exhibit a greater intertrial variability in their grip force application. We expected differences in these findings for the two force levels because the neural activity might be dependent on the magnitude of force (Dettmers et al., 1996; Ehrsson et al., 2001; Hendrix et al., 2009; Perez and Cohen, 2009; Parikh et al., 2014).

## MATERIALS AND METHODS

### Subjects

Thirteen young, healthy, right-handed subjects (Oldfield, 1971) aged between 18 and 36 years (mean ± SD: 25.30 ± 3.59 years; 4 females) provided written informed consent to participate in this study. Subjects eligible for the protocol had a normal or corrected-to-normal vision, no upper limb injury, and no history of neurological diseases or musculoskeletal disorders. They were screened for potential risks or adverse reactions to TMS using the TMS Adult Safety Screen questionnaire (Keel et al., 2001; Rossi et al., 2009). The study was approved by the Institutional Review Board of the University of Houston.

### Experimental apparatus

Subjects were instructed to grasp a custom-designed inverted T-shaped grip device using their index finger and thumb. Two six-dimensional force/torque transducers (Nano-25; ATI Industrial Automation, Garner, NC), mounted on the grip device, measured the force and moments exerted by their index finger and thumb on the object (Fig. 1). Force data were acquired using 12-bit analog-to-digital converter boards (sampling frequency 1 kHz; model PCI-6225; National Instruments, Austin, TX).

**Figure 1.**
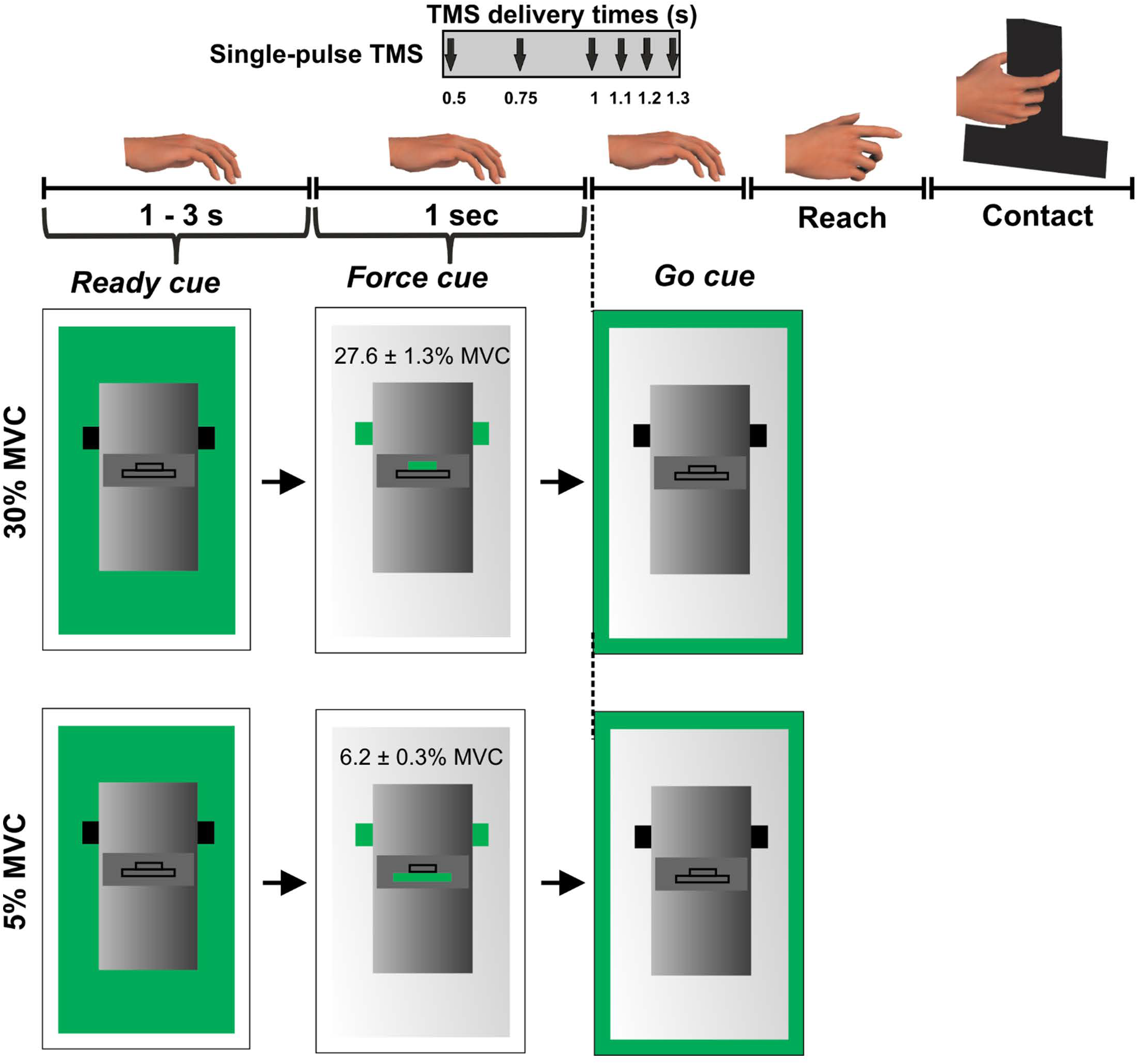
Experimental protocol. Figure adapted from *Parikh PJ, Davare M, McGurrin P, Santello M (2014) Corticospinal excitability underlying digit force planning for grasping in humans. Journal of Neurophysiology, 111: 2560–2569*.

### Electromyography (EMG)

We recorded muscle activity from first dorsal interosseous (FDI), abductor policis brevis (APB), and abductor digiti minimi (ADM) using differential surface electrodes (Delsys Bagnoli EMG System, Boston, MA). EMG data were sampled at 5 kHz using CED data acquisition board (Micro1401, Cambridge, England). Both force and EMG data were analysed using custom-made MATLAB script (R2016b; Mathworks, Natick, MA).

### Transcranial Magnetic Stimulation (TMS)

Single-pulse TMS was used to assess CSE during the experiment (Parikh et al., 2014; Davare et al., 2019). We first estimated the resting motor threshold (rMT) by delivering suprathreshold single monophasic TMS pulses (Magstim 200, Whitland, UK) with the TMS coil held tangential to the scalp and perpendicular to the presumed direction of the central sulcus, 45° from the midsagittal line, with the handle pointing backward, inducing current in the posteroanterior direction. The coil position was adjusted to optimize the motor-evoked potential (MEP) in all recorded muscles. Following this procedure, the rMT was estimated as the minimum TMS-intensity to elicit motor evoked potential (MEP) with an amplitude of ~50 µV (peak-to-peak) for at least 5 of the 10 consecutive trials in the FDI muscle (Klein-Flugge et al., 2013; Parikh et al., 2014; Rossini et al., 2015; Parikh and Santello, 2017). The TMS coil was stabilized using a coil holder mounted on the TMS chair (Rogue Research). The TMS coil was traced on the subject’s scalp using a surgical marker pen. The coil location was regularly checked for any displacement that might have occurred during a session. The average rMT across subjects (mean ± SE) was 41 ± 3% of the maximum stimulator output.

### Experimental design

Eleven of thirteen subjects participated in two experiments performed at least 24 hours apart. Two subjects were able to participate in one of the two experiments.

#### Experiment 1 (at rest; n = 12)

We established a relation between the variability in MEP and its amplitude at rest. We delivered single pulse TMS at the following TMS intensities: 0.9, 1, 1.1, 1.2, 1.3, 1.4, 1.5, 1.6 or 1.7 times the rMT (Darling et al., 2006) with ten consecutive pulses delivered at each intensity in a randomized order. Subjects neither performed a task nor received a stimulus except TMS.

#### Experiment 2 (the force task; n = 12)

During this session, we asked subjects to perform an isometric force production task using their index finger and thumb of the right hand. The distance between the grip device and the hand was ~30 cm at the beginning of each trial. Subjects were instructed to reach for the grip device, grasp the device at the same locations, and exert grip force to match a target on computer monitor using their index finger and thumb (Fig. 1).

We introduced two different force levels (30% and 5% of maximal pinch force) to investigate modulation in MEP variability at different force magnitudes. To rule out differential planning of digit position from trial-to-trial, we instructed subjects to grasp the device at marked locations on every trial. This location was denoted by a black tape attached on the front panel of the grip device (Parikh et al., 2014). A computer monitor placed behind the device displayed three sequential visual cues on every trial: ‘ready’, ‘force’ and ‘go’. The ‘ready’ cue signalled the beginning of a trial. The ‘force’ cue informed the subject about whether the upcoming force task required 5% or 30% of grip force application. Finally, the ‘go’ cue instructed subjects to initiate the reach and perform the force production task. The ‘ready’ and ‘force’ cues were separated by a randomly varying interval between 1-3s while ‘force’ and ‘go’ cues were separated by 1s (Fig. 1). Subjects were instructed to apply grip force to reach the target (displayed on the computer monitor during the ‘force’ cue presentation) at a self-selected speed and maintain that force for 3s using their right hand. Visual feedback of subject’s grip force was provided during each trial. Subjects practiced the force production task to get familiarized with the experimental task before the session. At the beginning of the session, we measured maximum voluntary contraction (MVC) for each subject by asking them to apply maximum pinch force only using right thumb and index finger. We selected the largest force of three MVC trials to set the force target.

While subjects performed the *force task*, single TMS pulses at 120% of rMT were delivered to the scalp location for FDI marked earlier at 1 of the 6 latencies in a randomized order: 0.5, 0.75, 1 (‘go’), 1.1, 1.2, or 1.3 s following the ‘force’ cue (Fig. 1). For this experimental setup, our earlier study has shown that the reach is not initiated at least until 0.4 s after the ‘go’ cue (i.e. 1.4 s following the ‘force’ cue) (Parikh et al., 2014). Therefore, the TMS pulse delivered at 6 different time points following the ‘force’ cue would represent planning related corticospinal activity. Each subject performed 15 trials per TMS time point and force level in a randomized sequence. As there were six TMS time points across two force levels, subjects performed 180 trials across four blocks with 45 trials per block and with ~5 min of rest between the blocks.

### Data processing and Statistical analysis

#### Behavioral variability in the application of force

We focused our analysis on the peak rate of force (PFR) application because it is known to be influenced by planning-related mechanisms (Johansson and Westling, 1988; Gordon et al., 1993). Specifically, we analyzed magnitude and time to peak force rate (Time_PFR_) to assess behavioral variability, as previously reported by (Flanagan and Beltzner, 2000; Poston et al., 2008). To compute the rate of grip force application, we first smoothed the grip force signal through a zero-phase lag, 4^th^ order, low-pass Butterworth filter (cutoff frequency: 14Hz) followed by calculating its first derivative with respect to the trial time (Flanagan and Beltzner, 2000). Separate analyses were then conducted for peak force rate (PFR) and Time_PFR_. Intertrial variability in these measures were assessed by calculating their standard deviation (SD) around the mean value for each TMS delivery time point and force level. We performed repeated measures analysis of variance (rmANOVA) with within-subject factors such as TMS (0.5, 0.75, 1, 1.1, 1.2, and 1.3 s) and FORCE (5% and 30% of force).

#### Intertrial variability in corticospinal excitability (CSE)

MEPs elicited using single TMS pulses were recorded to estimate the CSE (Parikh and Santello, 2017) during both sessions (Klein-Flugge et al., 2013; Parikh et al., 2014; Parikh and Santello, 2017). MEPs were also identified with pre-stimulus EMG contamination if the signal within 100 ms before TMS contained a peak-to-peak amplitude ≥0.1 mV (Klein-Flugge et al., 2013) and were removed from subsequent analysis (~2% of trials per subject). The data were divided into 9 bins (10 trials each) per subject representing MEPs at a given intensity (Klein-Flugge et al., 2013). Bins with more than 5 trials were included if the average MEP exceeded 0.1 mv (Klein-Flugge et al., 2013). Bins with average MEPs exceeding the average MEP amplitude from all the bins by three standard deviations were identified and excluded from further analysis (Klein-Flugge et al., 2013). For *experiment 2*, MEPs with pre-EMG contamination (peak-to-peak signal ≥0.1 mV within 100 ms before TMS pulse; ~2% trials per subject) and MEP measuring <0.1 mV were discarded from the subsequent analysis (~8% trials per subject), thus matching the criteria used for *experiment 1.* Removal of trials with background EMG assured that the TMS pulse delivered at different time points reflected planning rather than execution related activity. Overall, ~10% of trials were excluded during data processing across FDI, APB, and ADM muscles and time points. Coefficient of variation (CV = SD/mean) of MEP was used to quantify the intertrial variability in MEP (Klein-Flugge et al., 2013).

From *experiment 1*, we modelled a relation between MEP amplitude and its variability at rest using data points from all subjects (Darling et al., 2006; Klein-Flugge et al., 2013). The function characterizing this relationship was identified as the logarithmic curve of the form:

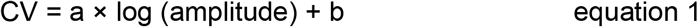

The coefficients *a* and *b* were identified using the *modelfun* function in MATLAB (Mathworks, Natick, MA). These coefficients were identified separately for three muscles (FDI, APB and ADM). We also assessed the residuals for the logarithmic fits for each muscle.

From *experiment 2*, Intertrial variability in MEPs (observed CV or CV_OBS_) was assessed by calculating the CV of MEPs (Klein-Flugge et al., 2013) for every TMS time point separately for each force level. The logarithmic model obtained from *experiment 1* was used to predict the CV of MEP (CV_PRED_) that was primarily due to intrinsic changes in MEP amplitude while preparing to perform the task. Such a model has been found to robustly estimate CV_PRED_ for any coefficient b and for the coefficient *a* within the range of −0.5 to infinity (Klein-Flugge et al., 2013). This predicted variability was then subtracted from the observed variability in CSE from *experiment 2* (CV_DIFF_ of MEP = CV_OBS_ – CV_PRED_). For CV_PRED_ and CV_DIFF_ of MEP, we performed separate rmANOVA (α = 0.05) with within-subject factors such as FORCE (2 levels: 5% and 30% of force) and TMS (6 levels: 0.5, 0.75, 1, 1.1, 1.2, and 1.3 s) for 3 muscles (FDI, APB, ADM).

To assess the repeatability of our findings, we performed this analysis on a dataset from 9 additional subjects. These subjects performed the force production task at either 10% of force or 1 N force under similar experimental paradigm (Parikh et al., 2014). As *experiment 1* was not conducted in the earlier study, we used the logarithmic model obtained using data points from all subjects in the current study. This logarithmic model resulted in the same results in *experiment 2*. These findings were presented earlier at the annual meeting of the Society for Neuroscience (Rao and Parikh, 2017) and are summarized in the Results section.

To assess the robustness of the analytical approach, we also analysed data without using a lower bound cut-off criterion for MEP amplitude and without using a bin-based cut-off criterion. Furthermore, the results may be sensitive to coefficients obtained by fitting a logarithmic model on data points from all subjects (i.e. a group-level model; *experiment 1* description above) versus fitting a separate model on data points from each subject (i.e. subject-level models). For each subject, a subject-level logarithmic model was used to calculate CV_PRED_ and the resulting CV_DIFF_ showed similar findings in *experiment 2* (see Results).

To investigate whether MEP variability during planning explained inter-individual differences in behavioural variability, we performed separate Pearson product-moment correlation analysis between the CV_DIFF_ of MEP and the behavioural measures i.e. SD of PFR and Time_PFR_ and CoP ellipse area.

#### EMG Analysis

We quantified the modulation in FDI and APB muscles involved in the production of grip force when subjects applied 30% and 5% of force on the object. For this purpose, we calculated the root mean square (RMS) value of the EMG signal for a 1.5s segment during steady force production at 5% and 30% of force separately for FDI and APB (Zhang et al., 2017; Hu et al., 2018). For data from each muscle, we performed paired t-test to compare the EMG activity measured at each force level.

#### Intertrial variability in digit placement

Although our experiment was designed to rule out differences in planning of digit position on each trial, it was important to confirm whether our design resulted in no difference in variability in digit contact points between 30% and 5% of force and between various TMS time points. To quantitatively evaluate this condition, we calculated the centre of pressure (CoP) for the thumb and index finger defined as the vertical and horizontal coordinates of the point of resultant force exerted by each digit on graspable surfaces of the grip device (Parikh and Cole, 2012).

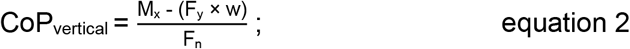

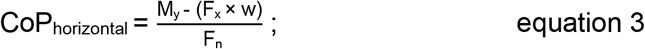

M_x_ and M_y_ are the moment about the x-axis and y-axis, respectively. F_x_ and F_y_ are the forces exerted on the grasp surface along the x-axis and y-axis, respectively. w is the distance between the surfaces of the F/T transducer and the grasp surface. F_n_ is the force component perpendicular to the grasp surface. To assess trial-to-trial variability in thumb and index finger CoPs, we computed area of an ellipse fitted to CoP (vertical and horizontal components calculated using equations 2 and 3). First, for each subject, we calculated an ellipse that contained CoP points within 95% confidence interval in each force level and at each TMS time point, separately for thumb and index finger. Surface area of these ellipses gave us a measure of intertrial variability in CoP across trials at a given TMS time point and at a given force level, as established in previous studies (Duarte and Zatsiorsky, 2002; Davare et al., 2007; Friendly et al., 2013). We performed separate rmANOVA with within-subject factors such as FORCE (2 levels: 30% and 5%) and TMS (6 levels: 0.5, 0.75, 1, 1.1, 1.2, and 1.3 s) for the thumb and index finger.

For all statistical analyses involving rmANOVA (α = 0.05), we used Mauchly’s test to assess the assumption of sphericity and applied Greenhouse-Geisser correction when needed. Post-hoc paired t-test comparisons were performed between adjacent TMS time points using Dunn-Sidak corrections. As stated earlier, we hypothesized that the modulation in CV of MEP and its relation with behavioural variability would be dependent on the magnitude of force (Dettmers et al., 1996; Ehrsson et al., 2001; Hendrix et al., 2009; Perez and Cohen, 2009; Parikh et al., 2014). Therefore, at each force level, we studied the modulation in CV of MEP using additional paired t-tests with appropriate Dunn-Sidak corrections and performed separate correlation analysis. The statistical analyses were performed using SPSS (version 25, SPSS Inc., Chicago, IL).

## RESULTS

### Intertrial variability in behavioral variables

#### Variability in time to peak force rate (Time_PFR_)

SD in Time_PFR_ from trial-to-trial was greater at 30% than 5% of force (main effect of FORCE: F_1,11_ = 31.160, p < 0.001, η_p_^2^ = 0.739; Fig. 2A). We observed no modulation in variability in Time_PFR_ across different TMS delivery time points for TMS pulses (neither a main effect of TMS: F_5,55_ = 1.370, p = 0.250, η_p_^2^ = 0.111, nor FORCE × TMS interaction: F_5,55_ = 0.469, p = 0.660, η_p_^2^ = 0.041).

**Figure 2.**
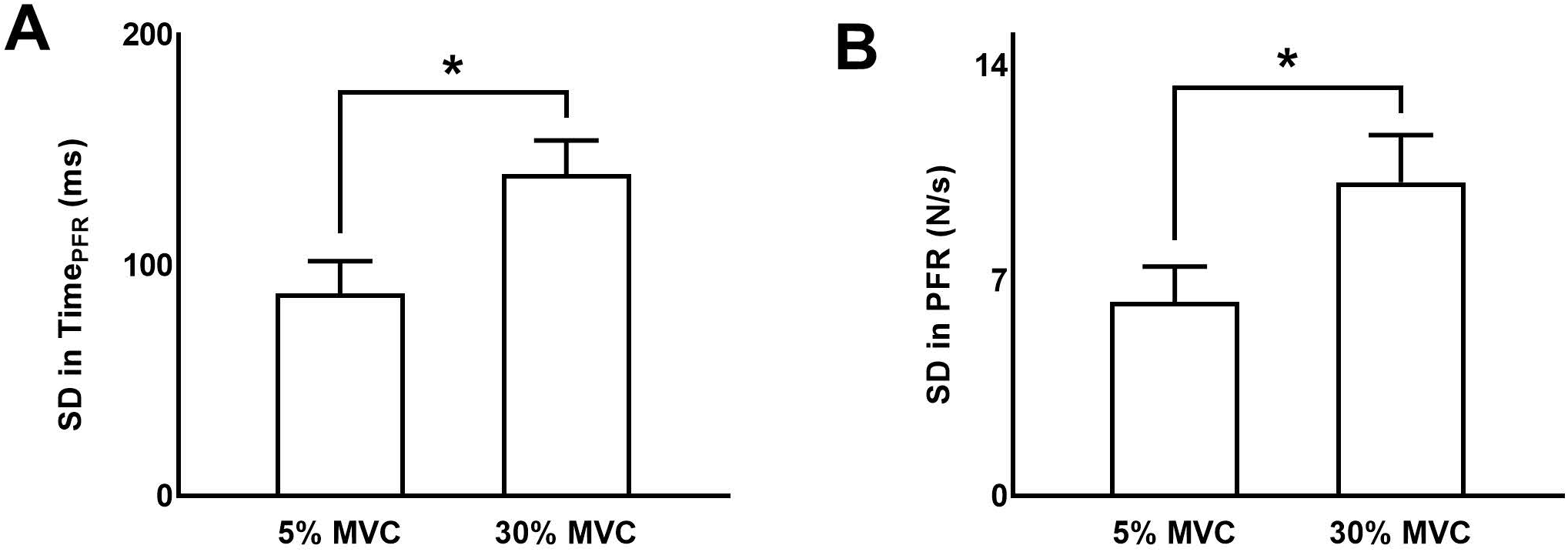
Behavioral variability. **A** and **B:** *S*tandard deviation (SD) in time to peak force rate and peak force rate, respectively, at 5% and 30% of force. Data are averages across all subjects (vertical bars denote SE). Asterisks indicate p < 0.05.

#### Variability in magnitude of peak force rate (PFR)

The standard deviation (SD) of PFR was greater at 30% than 5% of force (main effect of FORCE: F_1,11_ = 26.732, p < 0.001, η_p_^2^ = 0.708; Fig. 2B). The delivery of TMS pulses at different time points of planning did not influence the variability in PFR (neither a main effect of TMS: F_5,55_ = 0.244, p = 0.787, η_p_^2^ = 0.022, nor FORCE × TMS interaction: F_5,55_ = 2.456, p = 0.089, η_p_^2^ = 0.183).

#### Variability in digit contact points (CoP)

Variability in CoP was assessed by calculating the area of 95% confidence interval ellipse containing the digit contact points (Duarte and Zatsiorsky, 2002; Davare et al., 2007; Friendly et al., 2013). For the index finger contact point, we did not find difference in ellipse area between 30% and 5% of force across TMS time points (no main effect of FORCE: F_1,11_ = 0.38, p = 0.55, η_p_^2^ = 0.034; no main effect of TMS: F_5,55_ = 0.562, p = 0.728, η_p_^2^ = 0.049; no FORCE × TMS interaction: F_5,55_ = 0.606; p = 0.695, η_p_^2^ = 0.052; mean ± SE at 30% = 2.75 ± 0.19 cm^2^ and 5% = 2.71 ± 0.23 cm^2^; Fig. 3). Similarly, for the thumb contact point, we did not find difference in ellipse area between 30% and 5% trials across TMS time points (30% = 3.67 ± 0.41 cm^2^ and 5% = 3.75 ± 0.40 cm^2^ - no FORCE × TMS interaction: F_5,55_= 0.58; p = 0.71, η_p_^2^= 0.05; no main effect of FORCE: F_1,11_ = 0.11, p = 0.75, η_p_^2^ = 0.01; no main effect of TMS time points: F_5,55_ = 0.976, p = 0.44, η_p_^2^ = 0.08). These results suggest that the intertrial variability in digit contact points was similar across force levels and TMS time points.

**Figure 3.**
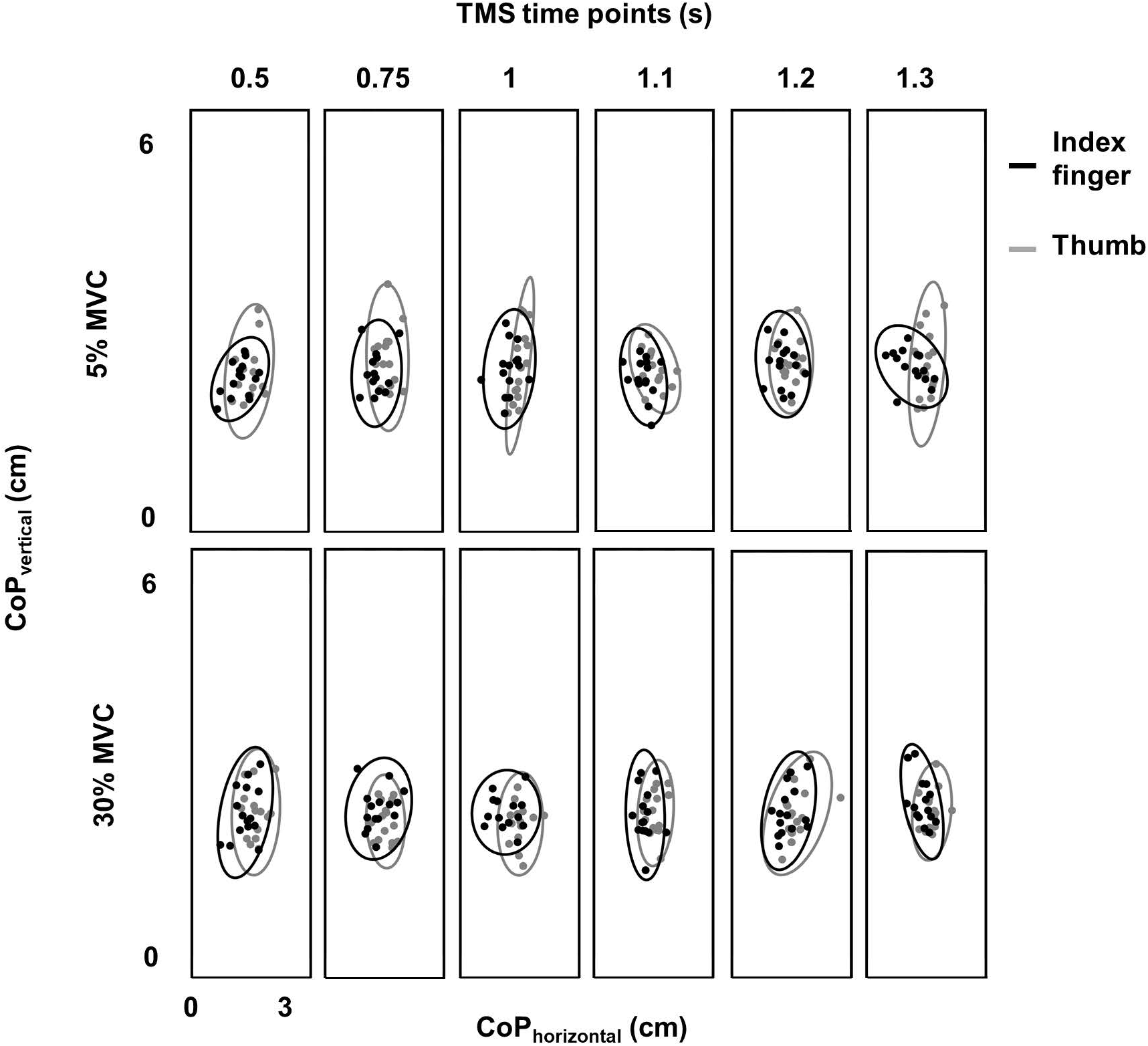
Variability in digit placement. Center of pressure (CoP) for thumb (gray) and index finger (black) for each TMS time point at 30% and 5% of force from a representative subject. Vertical and horizontal components of thumb and index finger CoP are shown on the same plot. Ellipse contained CoP points within 95% confidence interval in each task and at each TMS time point.

### Relation between MEP CV with MEP amplitude at rest

We modelled a relation between CV of MEP and amplitude of MEP separately for each muscle (FDI, APB and ADM). The logarithmic relationship, as described in equation 1, for FDI was as below:

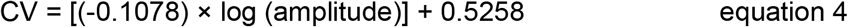

The values of the coefficients (*a*, *b*) from equation 1 for APB were (−0.0899, 0.4764) and for ADM were (−0.0773, 0.3116). The logarithmic fit and the residuals for FDI are shown in Fig. 4A.

**Figure 4.**
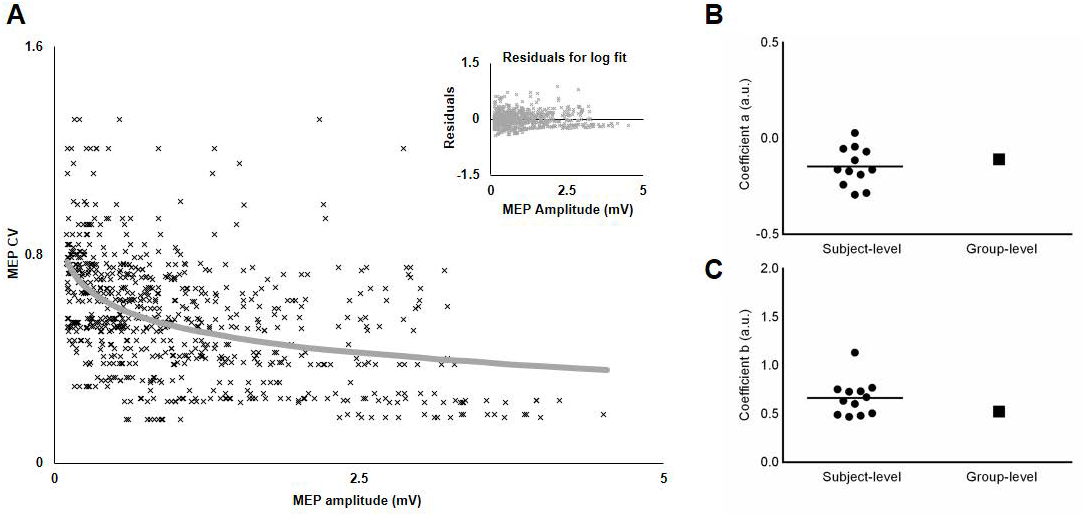
Relationship between variability (CV) and amplitude of MEP. **A.** The decrease in MEP CV with increase in MEP amplitude was characterized by a logarithmic fit. Inset plot shows trend in residuals for the logarithmic fit. **B.** Comparison of coefficient *a* between subject-level and group-level logarithmic models. **C.** Comparison of coefficient *b* between subject-level and group-level logarithmic models. Solid circles represent individual coefficients and the horizontal line represents mean coefficient.

### Variability in MEP due to changes in MEP amplitude (CV_PRED_) during task planning

We predicted MEP variability (CV_PRED_) while planning to exert isometric grip force for individual subjects using the logarithmic model separately for each muscle (equations 1 and 4). For FDI, we found that CV_PRED_ of MEP was different across TMS time points (main effect of TMS: F_5,55_ = 3.695, p = 0.006, η_p_^2^ = 0.251; Fig. 5A). However, this time-dependent modulation of CV_PRED_ of MEP was not different across force conditions (No FORCE × TMS interaction: F_5,55_ = 0.506, p = 0.770, η_p_^2^ = 0.044; no main effect of FORCE: F_1,11_ = 0.701, p = 0.420, η_p_^2^ = 0.060). Post hoc comparisons found a significant increase in CV_PRED_ of MEP from 0.5s to 0.75s (t_11_ = 3.2, p = 0.009, Cohen’s d_Z_ = 0.92). No difference in CV_PRED_ of MEP was found between other adjacent TMS time points (all p values > 0.05). Within each force level, we found significant increase in CV_PRED_ of MEP from 0.5s to 0.75s at 30% (t_11_ = 2.9, p = 0.015, Cohen’s d_Z_ = 0.84; Figs. 5A and 5B), but not at 5% of force (t_11_ = 2.1, p = 0.06, Cohen’s d_Z_ = 0.61). The change in CV_PRED_ of MEP at 30% of force was related to the change in MEP amplitude. Specifically, we found a decrease in MEP amplitude for FDI from 0.5s to 0.75s at 30% (t_11_ = 2.9, p = 0.014, Cohen’s d_Z_ = 0.84), but not at 5% (t_11_ = 1.4, p = 0.2, Cohen’s d_Z_ = 0.40) of force (Fig. 5C).

**Figure 5.**
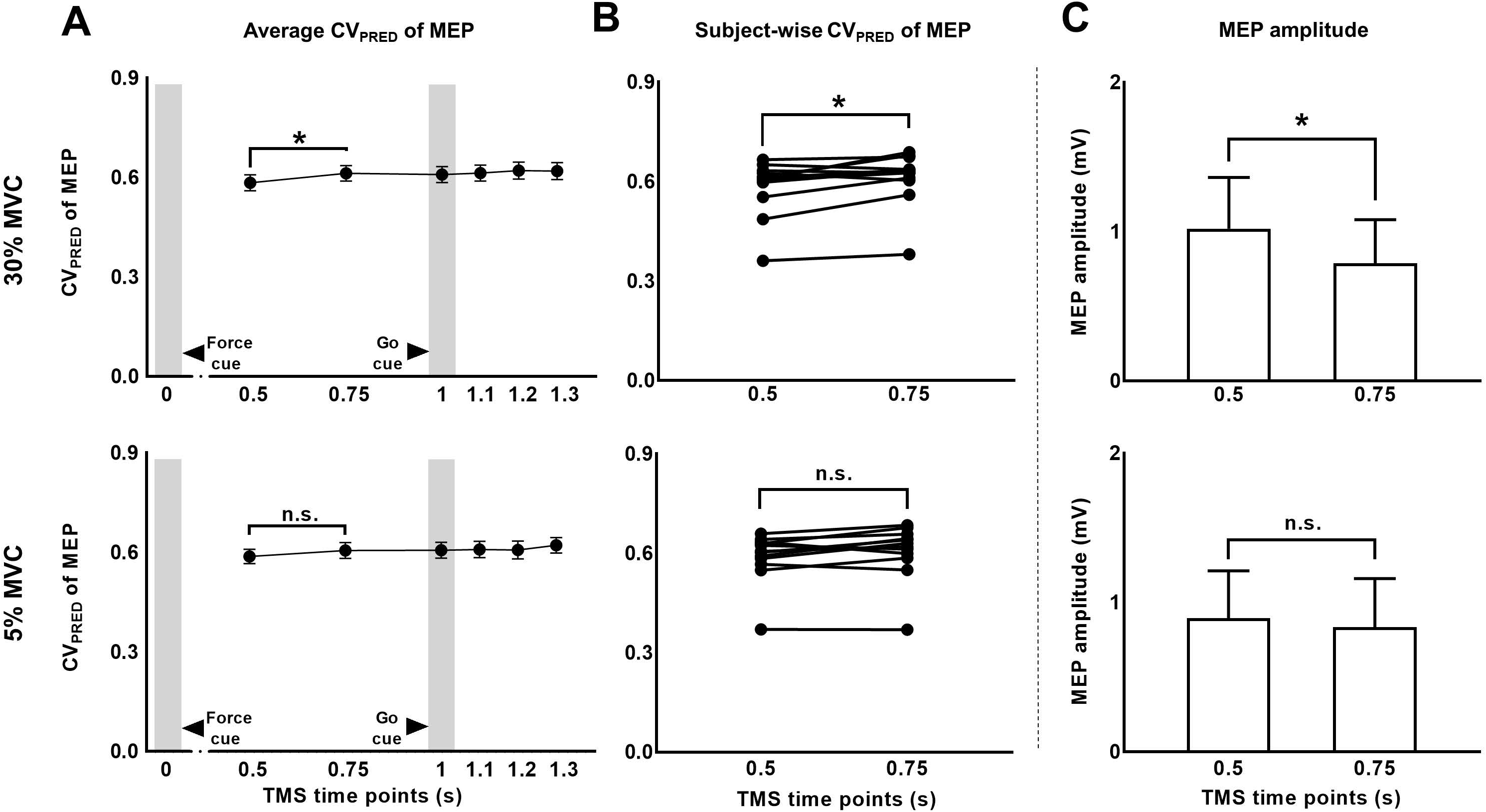
MEP CV due to changes in MEP amplitude during task preparation. **A.** Time-course of predicted CV of MEP at 30% compared to 5% of force. **B.** Subject-wise predicted CV of MEP data indicates a consistent rise across subjects from 0.5s and 0.75s at 30%, but not at 5% of force. **C.** A significant reduction in MEP amplitude from 0.5s to 0.75s explained the rise in predicted CV of MEP at 30% of force. Data in **A** and **C** are averages of all subjects (vertical bars denote SE). Asterisks indicate p < 0.016 and n.s. indicates p > 0.05.

For APB, we did not observe modulation in CV_PRED_ of MEP across force conditions and TMS time points (Table 1; No FORCE × TMS interaction: F_5,45_ = 0.588, p = 0.709, η_p_^2^ = 0.061; no main effect of FORCE: F_1,9_ = 2.313, p = 0.163, η_p_^2^ = 0.204; and no main effect of TMS: F_5,45_ = 0.988, p = 0.436, η_p_^2^ = 0.099). Similarly, for ADM, there was no modulation in predicted CV of MEP for across tasks and TMS time points (Table 1; No FORCE × TMS: F_5,35_ = 0.724, p = 0.610, η_p_^2^ = 0.094; no main effect of FORCE: F_1,7_ = 0.841, p = 0.390, η_p_^2^ = 0.107; and no main effect of TMS: F_5,35_ = 0.090, p = 0.993, η_p_^2^ = 0.013).

**Table 1:** Predicted CV (CV_PRED_) of MEP for each TMS Time Point and Force Level

| Predicted CV of MEP |  |  |  |  |  |  |  |  |  |  |  |  |
| --- | --- | --- | --- | --- | --- | --- | --- | --- | --- | --- | --- | --- |
| TMS<br>time<br>point (s) | FDI |  |  |  | APB |  |  |  | ADM |  |  |  |
|  | 5% MVC |  | 30% MVC |  | 5% MVC |  | 30% MVC |  | 5% MVC |  | 30% MVC |  |
|  | Mean | SD | Mean | SD | Mean | SD | Mean | SD | Mean | SD | Mean | SD |
| 0.5 | 0.588 | 0.076 | 0.584 | 0.084 | 0.506 | 0.055 | 0.506 | 0.063 | 0.416 | 0.058 | 0.401 | 0.059 |
| 0.75 | 0.606 | 0.084 | 0.613 | 0.081 | 0.526 | 0.054 | 0.512 | 0.043 | 0.415 | 0.057 | 0.406 | 0.056 |
| 1 | 0.607 | 0.085 | 0.609 | 0.085 | 0.528 | 0.056 | 0.521 | 0.070 | 0.406 | 0.066 | 0.408 | 0.057 |
| 1.1 | 0.609 | 0.087 | 0.613 | 0.085 | 0.511 | 0.049 | 0.523 | 0.065 | 0.406 | 0.058 | 0.403 | 0.064 |
| 1.2 | 0.607 | 0.093 | 0.621 | 0.088 | 0.513 | 0.055 | 0.504 | 0.044 | 0.411 | 0.057 | 0.406 | 0.060 |
| 1.3 | 0.621 | 0.082 | 0.619 | 0.088 | 0.512 | 0.062 | 0.512 | 0.058 | 0.411 | 0.064 | 0.408 | 0.059 |

### MEP variability above and beyond predicted CV of MEP during task planning (CV_DIFF_)

To investigate whether MEP variability modulated beyond CV_PRED_ of MEP while planning for the force task, we subtracted CV_PRED_ of MEP (Table 1) from CV_OBS_ of MEP (Table 2) to obtain CV_DIFF_ of MEP (Table 3). For FDI, we found modulation in CV_DIFF_ of MEP across TMS time points (main effect of TMS: F_5,55_ = 4.730, p = 0.001, η_p_^2^ = 0.301). However, CV_DIFF_ of MEP was similar across force conditions (no FORCE × TMS interaction: F_5,55_ = 0.436, p = 0.821, η_p_^2^ = 0.038 and no main effect of FORCE: F_1,11_ = 0.065, p = 0.803, η_p_^2^ = 0.006). Post hoc comparisons found a significant increase in CV_DIFF_ of MEP from 1.2s to 1.3s (t_11_ = 3.1, p = 0.01, Cohen’s d_Z_ = 0.89). No difference in CV_DIFF_ of MEP was found between other adjacent TMS time points (all p values > 0.13). Within each force level, we found significant increase in CV_DIFF_ of MEP from 1.2s to 1.3s at 30% (t_11_ = 2.9, p = 0.015, Cohen’s d_Z_ = 0.84; Fig. 6A) but not at 5% (t_11_ = 1.6, p = 0.14, Cohen’s d_Z_ = 0.46) of force. Most subjects showed a systematic increase in CV_DIFF_ of MEP from 1.2s compared with 1.3s at 30% of force (9 of 12 subjects; Fig. 6B). However, at 5% of force, the change in CV_DIFF_ of MEP from 1.2s to 1.3s was not consistent across subjects. Although the variability related to MEP amplitude were removed to obtain CV_DIFF_ of MEP, we confirmed that there was no change in MEP amplitude from 1.2s to 1.3s (30%: t_11_ = 0.76, p = 0.47, Cohen’s d_Z_ = 0.22; 5%: t_11_ = 1.04, p = 0.32, Cohen’s d_Z_ = 0.30; Fig. 6C).

**Table 2:** Observed CV (CV_OBS_) of MEP for each TMS Time Point and Force Level

| Observed CV of MEP |  |  |  |  |  |  |  |  |  |  |  |  |
| --- | --- | --- | --- | --- | --- | --- | --- | --- | --- | --- | --- | --- |
| TMS<br>time<br>point (s) | FDI |  |  |  | APB |  |  |  | ADM |  |  |  |
|  | 5% MVC |  | 30% MVC |  | 5% MVC |  | 30% MVC |  | 5% MVC |  | 30% MVC |  |
|  | Mean | SD | Mean | SD | Mean | SD | Mean | SD | Mean | SD | Mean | SD |
| 0.5 | 0.585 | 0.177 | 0.705 | 0.256 | 0.540 | 0.195 | 0.546 | 0.187 | 0.395 | 0.158 | 0.378 | 0.210 |
| 0.75 | 0.681 | 0.249 | 0.683 | 0.273 | 0.628 | 0.281 | 0.706 | 0.276 | 0.500 | 0.322 | 0.495 | 0.331 |
| 1 | 0.635 | 0.163 | 0.618 | 0.174 | 0.577 | 0.222 | 0.499 | 0.211 | 0.421 | 0.189 | 0.594 | 0.287 |
| 1.1 | 0.586 | 0.184 | 0.551 | 0.197 | 0.617 | 0.273 | 0.523 | 0.246 | 0.478 | 0.176 | 0.428 | 0.191 |
| 1.2 | 0.589 | 0.202 | 0.606 | 0.206 | 0.537 | 0.184 | 0.551 | 0.206 | 0.472 | 0.296 | 0.462 | 0.274 |
| 1.3 | 0.799 | 0.345 | 0.789 | 0.327 | 0.529 | 0.194 | 0.548 | 0.189 | 0.473 | 0.296 | 0.459 | 0.185 |

**Table 3:**
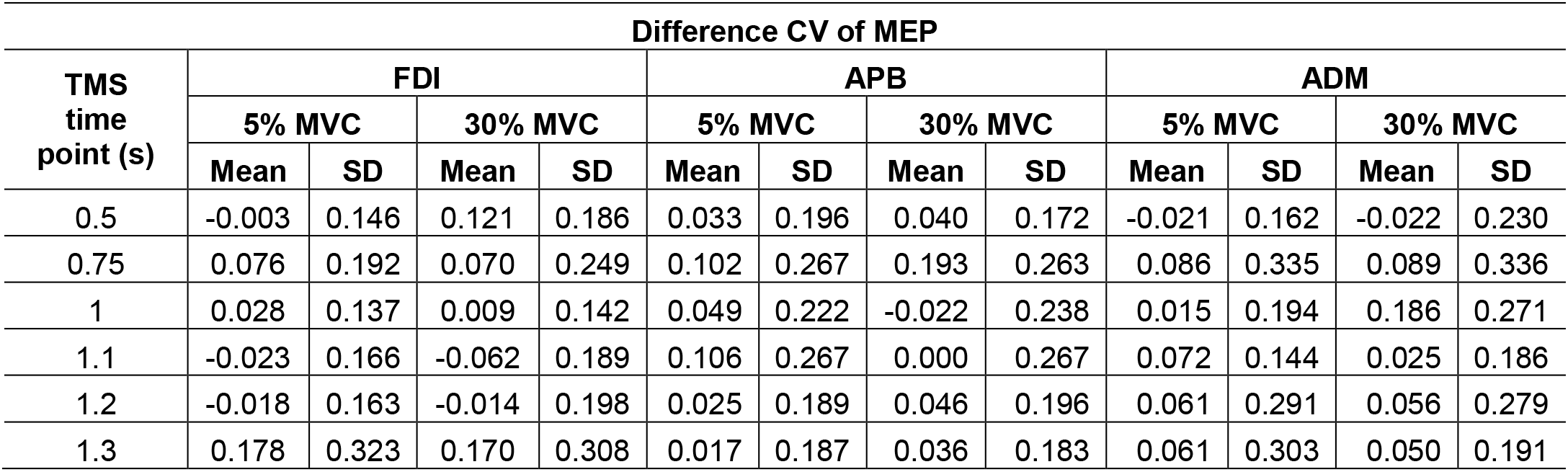
Difference CV (CV_DIFF_) of MEP for each TMS Time Point and Force Level

| Difference CV of MEP |  |  |  |  |  |  |  |  |  |  |  |  |
| --- | --- | --- | --- | --- | --- | --- | --- | --- | --- | --- | --- | --- |
| TMS<br>time<br>point (s) | FDI |  |  |  | APB |  |  |  | ADM |  |  |  |
|  | 5% MVC |  | 30% MVC |  | 5% MVC |  | 30% MVC |  | 5% MVC |  | 30% MVC |  |
|  | Mean | SD | Mean | SD | Mean | SD | Mean | SD | Mean | SD | Mean | SD |
| 0.5 | -0.003 | 0.146 | 0.121 | 0.186 | 0.033 | 0.196 | 0.040 | 0.172 | -0.021 | 0.162 | -0.022 | 0.230 |
| 0.75 | 0.076 | 0.192 | 0.070 | 0.249 | 0.102 | 0.267 | 0.193 | 0.263 | 0.086 | 0.335 | 0.089 | 0.336 |
| 1 | 0.028 | 0.137 | 0.009 | 0.142 | 0.049 | 0.222 | -0.022 | 0.238 | 0.015 | 0.194 | 0.186 | 0.271 |
| 1.1 | -0.023 | 0.166 | -0.062 | 0.189 | 0.106 | 0.267 | 0.000 | 0.267 | 0.072 | 0.144 | 0.025 | 0.186 |
| 1.2 | -0.018 | 0.163 | -0.014 | 0.198 | 0.025 | 0.189 | 0.046 | 0.196 | 0.061 | 0.291 | 0.056 | 0.279 |
| 1.3 | 0.178 | 0.323 | 0.170 | 0.308 | 0.017 | 0.187 | 0.036 | 0.183 | 0.061 | 0.303 | 0.050 | 0.191 |

**Figure 6.**
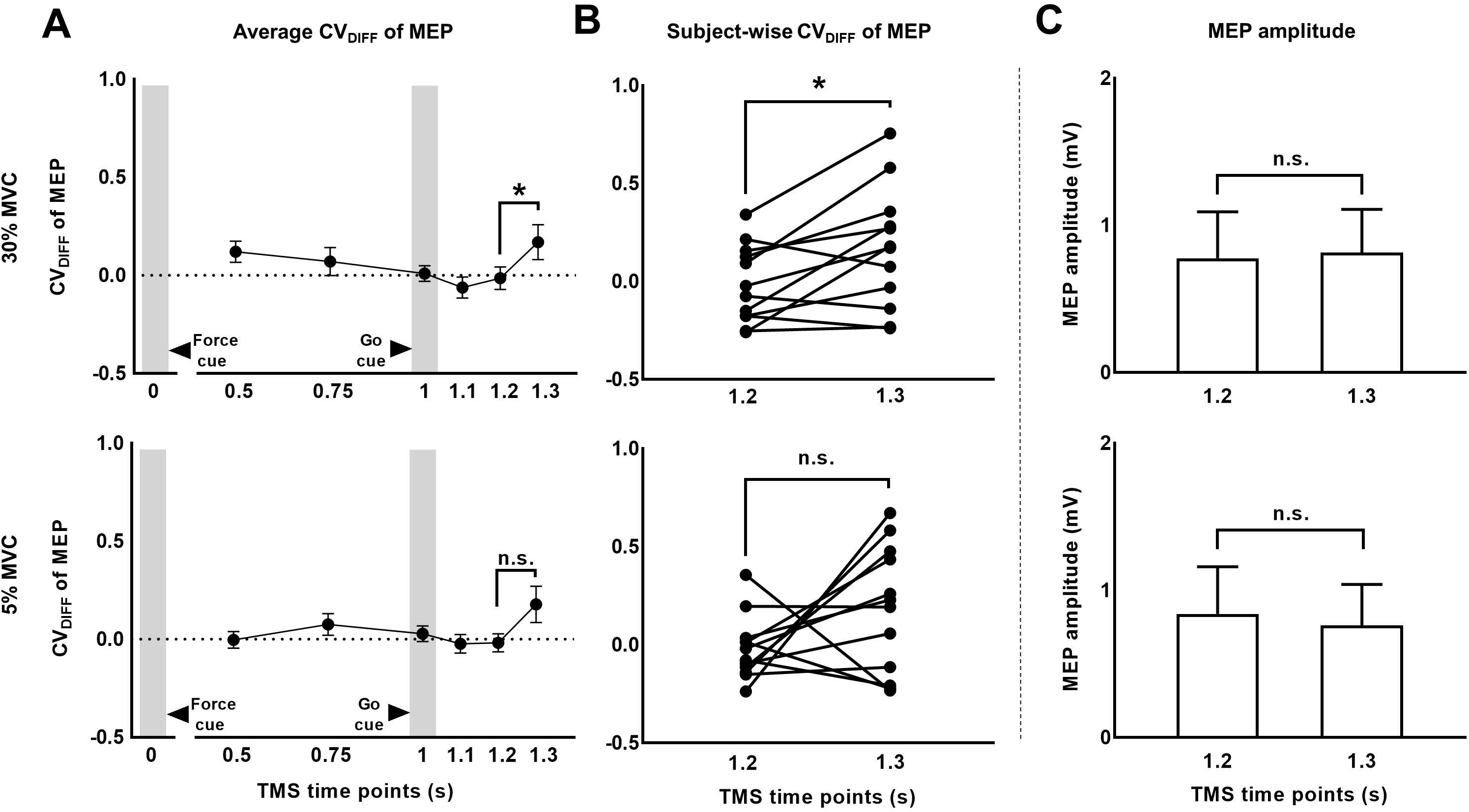
MEP CV rose above and beyond changes in MEP amplitude. **A.** Time-course of CV_DIFF_ (= observed – predicted CV) of MEP at 30% and 5% of force. **B.** Subject-wise CV of MEP data indicates a consistent rise across subjects from 1.2s and 1.3s at 30%, but not at 5% of force. **(C)** MEP amplitude analysis showed no modulation from 1.2s to 1.3s. Data in **A** and **C** are averages of all subjects (vertical bars denote SE). Asterisks indicate p < 0.016 and n.s. indicates p>0.1.

For APB, CV_DIFF_ of MEP was not different across force conditions and TMS time points (Table 3; no FORCE × TMS time points interaction: F_5,45_ = 0.302, p = 0.909, η_p_^2^ = 0.032; no main effect of TMS: F_5,45_ = 1.953, p = 0.104, η_p_^2^ = 0.178, and no main effect of Force: F_1,9_ = 0.290, p = 0.603, η_p_^2^ = 0.031). Similarly, for ADM, CV_DIFF_ of MEP was not different across force conditions and TMS time points (Table 3; no FORCE × TMS interaction: F_5,35_ = 0.746, p = 0.532, η_p_^2^ = 0.096; no main effect of TMS: F_5,35_ = 2.880, p = 0.073, η_p_^2^ = 0.118, and no main effect of FORCE: F_1,7_ = 0.938, p = 0.365, η_p_^2^ = 0.118).

To understand muscle-specific modulation in CV_PRED_ and CV_DIFF_ of MEP, we investigated modulation in FDI and APB EMG activity at 30% and 5% of force. We found that the EMG activity was greater for 30% versus 5% of force for FDI (t_11_ = 2.7, p = 0.019, Cohen’s d_Z_ = 0.78), but not for APB (t_11_ = 1.4, p = 0.18, Cohen’s d_Z_ = 0.40; Fig. 7), thus suggesting asymmetrical contribution of FDI and APB in the application of grip force, in agreement with previous reports (Li et al., 2013, 2015; Nataraj et al., 2015).

**Figure 7.**
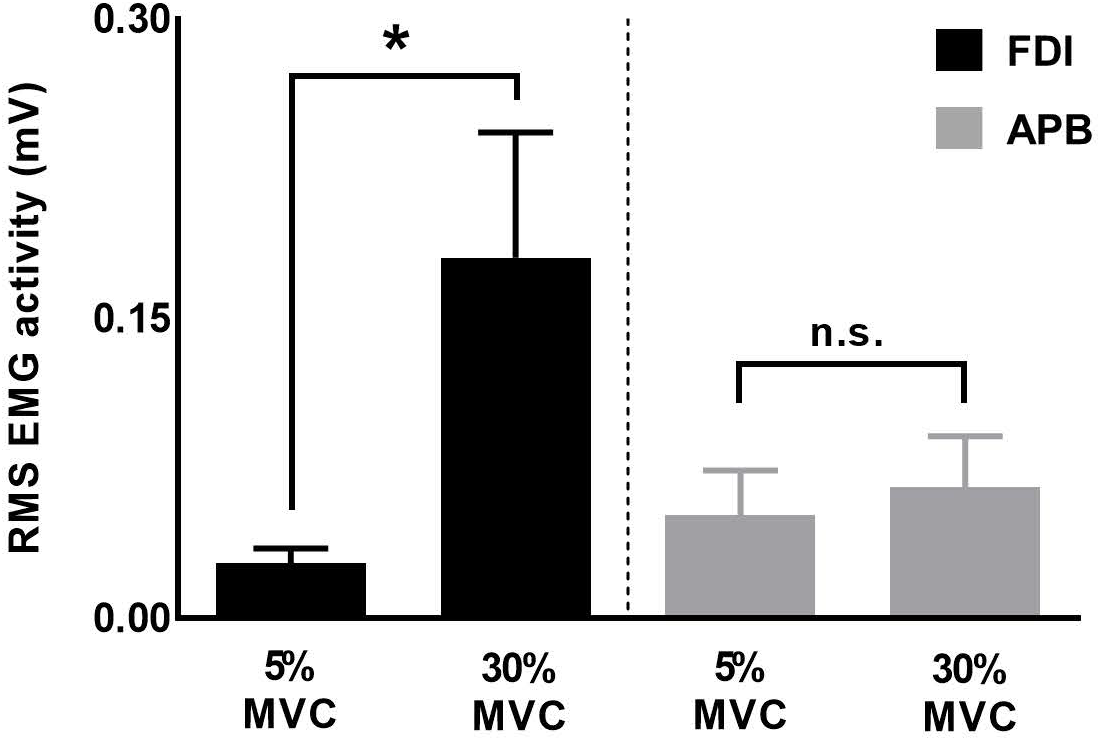
EMG activity for FDI and APB muscles. Force magnitude-dependent modulation in EMG activity was significant for FDI but not for APB muscles. Data are averages of all subjects (vertical bars denote SE), asterisk indicates p = 0.019 and n.s. indicates p>0.1.

### Correlation between the rise in CV of MEP during planning and behavioral variability

We investigated whether the increase in task-related MEP variability (i.e. CV_DIFF_ of MEP) from 1.2s to 1.3s explained the inter-individual differences in trial-to-trial behavioral variability. We found that the increase in CV_DIFF_ of MEP from 1.2s to 1.3s explained 64% of inter-individual differences in Time_PFR_ SD (Pearson’s *r* = 0.80, p = 0.0017; Fig. 8) at 30% of force. However, similar association between CV_DIFF_ of MEP and SD in Time_PFR_ was not observed for 5% of force (r = −0.25, p = 0.42). We also found no correlation between CV_DIFF_ of MEP and SD of PFR or CoP variability (all r-values < 0.26, all p values > 0.42).

**Figure 8.**
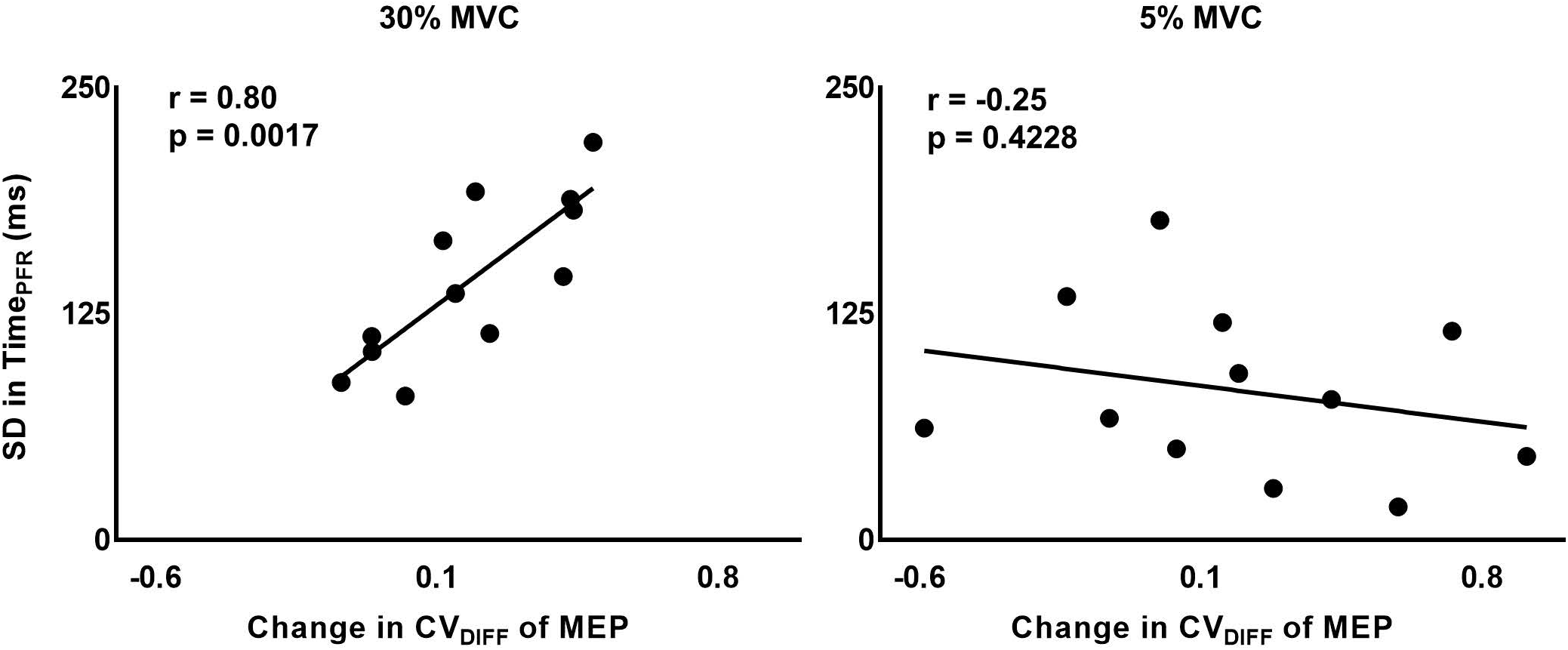
Correlation between intertrial task-specific variability in MEP and time to peak force rate. Modulation in intertrial MEP variability (CV_DIFF_ of MEP) for FDI muscle explained inter-individual differences in the trial-to-trial fluctuations in time to peak force rate, selectively at 30% (r = 0.80, p = 0.0017), but not at 5% (r = −0.25, p = 0.4228) of force.

### Robustness and Repeatability of our findings

To test the robustness of the findings with respect to MEP pre-processing, we analyzed our data using no lower bound cut-off for MEP amplitude and no bin-based cut-off criteria (see *Methods*). Furthermore, we fitted a logarithmic model for each subject’s data points to understand if the findings were sensitive to the group-level model (see *Methods*). The results were similar to that reported above. Across individual logarithmic models, the coefficient *a* ranged from −0.29 to 0.029 and the coefficient *b* ranged from 0.47 to 1.13 (Fig. 4B and 4C).

The logarithmic model obtained from *experiment 1* was used to obtain CV_PRED_ for each subject. As done earlier, we subtracted CV_PRED_ of MEP from CV_OBS_ of MEP to obtain CV_DIFF_ of MEP. We found modulation in CV_DIFF_ of MEP in FDI across TMS time points (main effect of TMS: F_5,55_ = 7.64, p < 0.001, η_p_^2^ = 0.41). CV_DIFF_ of MEP was similar across force conditions (no FORCE × TMS interaction: F_5,55_ = 0.55, p = 0.73, η_p_^2^ = 0.048 and no main effect of FORCE: F_1,11_ = 0.021, p = 0.88, η_p_^2^ = 0.002). Post hoc comparisons found a significant increase in CV of MEP from 1.2s to 1.3s (t_11_ = 3.92, p = 0.002, Cohen’s d_Z_ = 1.13). No difference in CV_DIFF_ of MEP was found between other adjacent TMS time points (all p values > 0.10). Within each force level, we found significant increase in CV_DIFF_ of MEP from 1.2s to 1.3s at 30% (t_11_ = 2.9, p = 0.015, Cohen’s d_Z_ = 0.84) but not at 5% (t_11_ = 1.9, p = 0.08, Cohen’s d_Z_ = 0.55) of force. Importantly, the relationship between the modulation in CV_DIFF_ of MEP and intertrial behavioral variability was preserved. That is, the increase in CV_DIFF_ of MEP from 1.2s to 1.3s explained 61% of inter-individual differences in Time_PFR_ SD (Pearson’s *r* = 0.77, p = 0.0029) at 30% of force.

To test the repeatability of the MEP findings, we separately analyzed data from 9 additional subjects who had performed a similar task (LF = 1N and HF = 10% of MVC) as described in (Parikh et al., 2014). We found modulation in CV_DIFF_ of MEP in FDI across TMS time points (main effect of TMS: F_5,40_ = 3.63, p = 0.0081, η_p_^2^ = 0.31). CV_DIFF_ of MEP was similar across force conditions (no FORCE × TMS interaction: F_5,40_ = 0.41, p = 0.84, η_p_^2^ = 0.048 and no main effect of FORCE: F_1,8_ = 0.029, p = 0.86, η_p_^2^ = 0.004). Post hoc comparisons found an increase in CV_DIFF_ of MEP from 1.2s to 1.3s (t_8_ = 2.61, p = 0.03, Cohen’s dz = 0.87), however it failed to reach the corrected significance level. No difference in CV_DIFF_ of MEP was found between other adjacent TMS time points (all p values > 0.15). Within each force level, we found an increase in CV_DIFF_ of MEP, although non-significant, from 1.2s to 1.3s at 30% of force (t_8_ = 2.16, p = 0.06, Cohen’s d_Z_ = 0.72). At 5% of force, we did not observe an increase in CV_DIFF_ of MEP from 1.2s to 1.3s (t_8_ = 1.8, p = 0.09, Cohen’s d_Z_ = 0.60) of force.

## DISCUSSION

We found that CSE variability increased beyond changes observed in CSE magnitude (i.e. CV_DIFF_ of MEP) while preparing to exert digit forces in a self-paced reach-to-grasp paradigm. The increase in CSE variability occurred after the ‘go’ cue presentation and this effect was temporally dissociated from the decrease in CSE magnitude that occurred before the ‘go’ cue presentation. The time-dependent modulation in CSE variability and CSE amplitude was evident at 30%, but not at 5% of force. Importantly, at 30% of force, individuals with larger increase in CSE variability also exhibited larger intertrial variability in time to peak force rate. These results were found to be repeatable across studies and robust to different data analysis methods. We discuss our findings in relation to potential sources underlying the increase in CSE variability and its contribution to the application of grip force.

### Modulation in CSE variability during task planning

Using a logarithmic model relating CSE magnitude and variability, we predicted the component of variability in CSE during task preparation that can be attributed to changes in CSE magnitude. We found a significant increase in predicted CV of MEP at 30%, but not at 5% of force. As predicted CV is primarily influenced by CSE magnitude, we found a corresponding reduction in CSE magnitude from 0.5s to 0.75s following the ‘force’ cue presentation at 30%, but not at 5% of force. This finding is consistent with our previous report demonstrating modulation in CSE magnitude at a higher force (Parikh et al., 2014). We further show that the intertrial variability in CSE rose beyond predicted variability in CSE at 30%, but not at 5% of force. Interestingly, the decrease in CSE magnitude and the increase in task-specific variability in CSE were temporally dissociated because the later occurred from 1.2s to 1.3s following the presentation of ‘force’ cue during task preparation. This finding suggests distinct neural sources underlying the modulation in CSE magnitude and the component of CSE variability not related to changes in its magnitude. Moreover, a consistent change in these variables across individuals at 30% of force (Figs. 5B and 6B) might represent important characteristics of individuals and thus the modulation in neural underpinnings during task planning (Kanai and Rees, 2011).

### Potential mechanisms that increased CSE variability during task planning

Increase in attentional demands may not explain the increase in CSE variability as it is usually associated with a reduction in neuronal firing rate variability (Cohen et al., 1997; Masquelier, 2013). Moreover, our experimental design ruled out any difference in planning of digit position from trial-to-trial between force levels and across time points. These findings suggest that the modulation in CSE variability was specific to grasp force planning during the reach-to-grasp task and provides information about fluctuations in planning-related corticospinal activity.

CSE arises from activation of intracortical circuitry within M1, cortico-cortical inputs to M1, and subcortical and spinal structures (Bestmann and Krakauer, 2015). The modulation in neuronal activity within primate M1 and premotor cortices has been found to depend on the magnitude of grasp force (Hendrix et al., 2009). Parietal, occipital, cerebellum, dorsolateral prefrontal cortex, and basal ganglia are also known to contribute to the planning of grip force (Dettmers et al., 1996; Ehrsson et al., 2000, 2001; Chouinard et al., 2005; Berner et al., 2007; Davare et al., 2007; Dafotakis et al., 2008). Virtual lesion studies using TMS have demonstrated contribution of human somatosensory, premotor dorsal, and supplementary motor regions in regulating the timing of digit force application (Davare et al., 2006; Schabrun et al., 2008; White et al., 2013). Evidence also exists in humans about the functional role of reticulospinal tracts in the control of coordinated hand movements such as those performed in our study (Honeycutt et al., 2013). It is less likely that changes in spinal motor neuron pool directly contributed to the increase in CSE variability during motor planning because variability in spinal motor neuronal excitability (as assessed by modulation in H-reflex) has been suggested to arise from changes in descending drive from supraspinal structures to spinal cord during motor planning (Collins et al., 1993; Misiaszek, 2003). Taken together, the increase in CSE variability observed during task preparation in our study is potentially sourced within supraspinal structures. Modulation in activation of these potential sources might have contributed to intertrial fluctuations in presynaptic inputs to M1 neurons (Lemon, 2008), thus resulting in modulation in CSE variability. As noted above, the inputs to M1 that influence CSE magnitude (Parikh et al., 2014) might be distinct from the inputs to M1 that influence CSE variability.

### Rise in CSE variability explains inter-individual differences in behavioral variability

In monkeys, neuronal firing rate variability during movement preparation has been suggested to explain ~50% of variability in reach speed from trial-to-trial (Churchland et al., 2006b, 2006a). Consistent with this primate work, we found that the intertrial variability in CSE specific to task-planning explained ~64% of inter-individual differences in behavioral, viz. Time_PFR_, variability in humans. The rise in CSE variability was associated with Time_PFR_ variability but not with variability in magnitude of peak force rate, although both factors are known to be important for accurate force application (Poston et al., 2008). It is possible that the intertrial variability in CSE during force planning may encode the variability in timing of force application as a control variable. To the best of our knowledge, this association provides first evidence in humans showing contribution of variability in planning-related neural mechanisms to motor output variability and corroborates earlier behavioral work in humans (van Beers, 2009). Fluctuations in neural activity during task execution may explain the remaining inter-individual differences in timing variability. A recent neuroimaging study found that the variability in BOLD-activity within intraparietal cortex recorded concurrently with task performance (i.e. during movement execution) accounts for ~25% of inter-individual differences in movement extent variability (Haar et al., 2017). In our study, the neural activity engaged during force planning may also be present during force execution and thus potentially contributing to the inter-individual differences in behavioral variability.

Overall, our study provides a novel insight into the contribution of planning related mechanisms to behavioral variability by assessing variability in human CSE in a self-paced reach-to-grasp paradigm. This is the first evidence showing that individuals with higher variability in neural activity during planning also exhibited a greater behavioral variability.

## ACKNOWLEDGEMENT

We thank Drs. Sheng Li, Stacey Gorniak, Marco Santello, and Joseph Francis for their comments on an earlier version of this manuscript.

